# Cold-sensing TRP channels and temperature preference modulate ovarian development in the model organism *Drosophila melanogaster*

**DOI:** 10.1101/2025.03.20.644370

**Authors:** Gabriele Andreatta, Sara Montagnese, Rodolfo Costa

## Abstract

Temperature is perceived primarily via Transient Receptor Potential (TRP) channels, which are integral to the molecular machinery sensing environmental and cellular signals. Functional evidence of TRP channels involvement in regulating cold-induced developmental/reproductive responses remains scarce. Here we show that mutations affecting cold-sensing TRP channels antagonize the reduction of ovarian development induced by low temperatures (reproductive dormancy) in *Drosophila melanogaster*. More specifically, mutants for *brv1, trp*, and *trpl* significantly lowered dormancy levels at 12°C, and exhibited well-developed oocytes characterized by advanced vitellogenesis. Similarly, functional knockouts for *norpA*, a gene encoding a phospholipase C acting downstream to Trp and Trpl, exhibited a reduced dormancy response, suggesting that Ca^2+^ signalling is key to relaying cold-sensing stimuli during dormancy induction and maintenance. Finally, mutants with altered temperature preference (i.e. exhibiting impaired cold or warm avoidance) differentially responded to cold, either lowering or increasing dormancy levels. In summary, our phenotypic analysis provides functional evidence of developmental/reproductive modulation by specific cold-sensing TRP channels in *Drosophila melanogaster*, and indicates that temperature preference affects developmental processes. As the studied genes are highly conserved and have mammalian homologues, the potential implications of our findings for human metabolism and drug development are also discussed.

## 1. Introduction

Despite the evolutionary distance, the fruitfly *Drosophila melanogaster* represents a powerful model organism to investigate aspects of human physiology and human genetic diseases, as several pathways involved are highly conserved [1–3]. Indeed, *Drosophila* research contributed to shed light on the pathophysiology of some forms of cancer [4,5], some neurological and metabolic disorders [6–10], mitochondrial disease [11,12], and some aspects of aging [13,14]. The fruitfly has played a significant role in the understanding of the molecular and cellular basis of temperature sensing [15,16]. This heavily relies on TRP channels, which were first discovered in *Drosophila* [17,18], and have also shown promise in drug development and translational/clinical applications [19,20].

The perception of environmental temperature modulates behaviour, metabolism and development in animals, favouring survival [21–24]. Over the past decades, progress has been made in defining the neuronal and molecular networks underlying thermosensation in different model organisms [21,25]. These studies show how temperature sensing relies extensively on Transient Receptor Potential (TRP) channels, a conserved protein group which mediates a wide array of sensory processes, including photoreception, hygrosensation, hearing, mechanosensation and nociception [26–28]. The TRP channel superfamily includes six families in bilatarians [19,29] (see Figure 1 and Supplementary Figure S1). Interestingly, in both flies and mammals, different classes of TRP channels are responsible for the perception of warm and cold temperature, activating different neuronal networks [26]. More specifically, cold sensing triggers an array of species-specific responses including cool avoidance, thermogenesis, as well as metabolic and developmental slowing/arrest [15,30,31].

**Figure 1.**
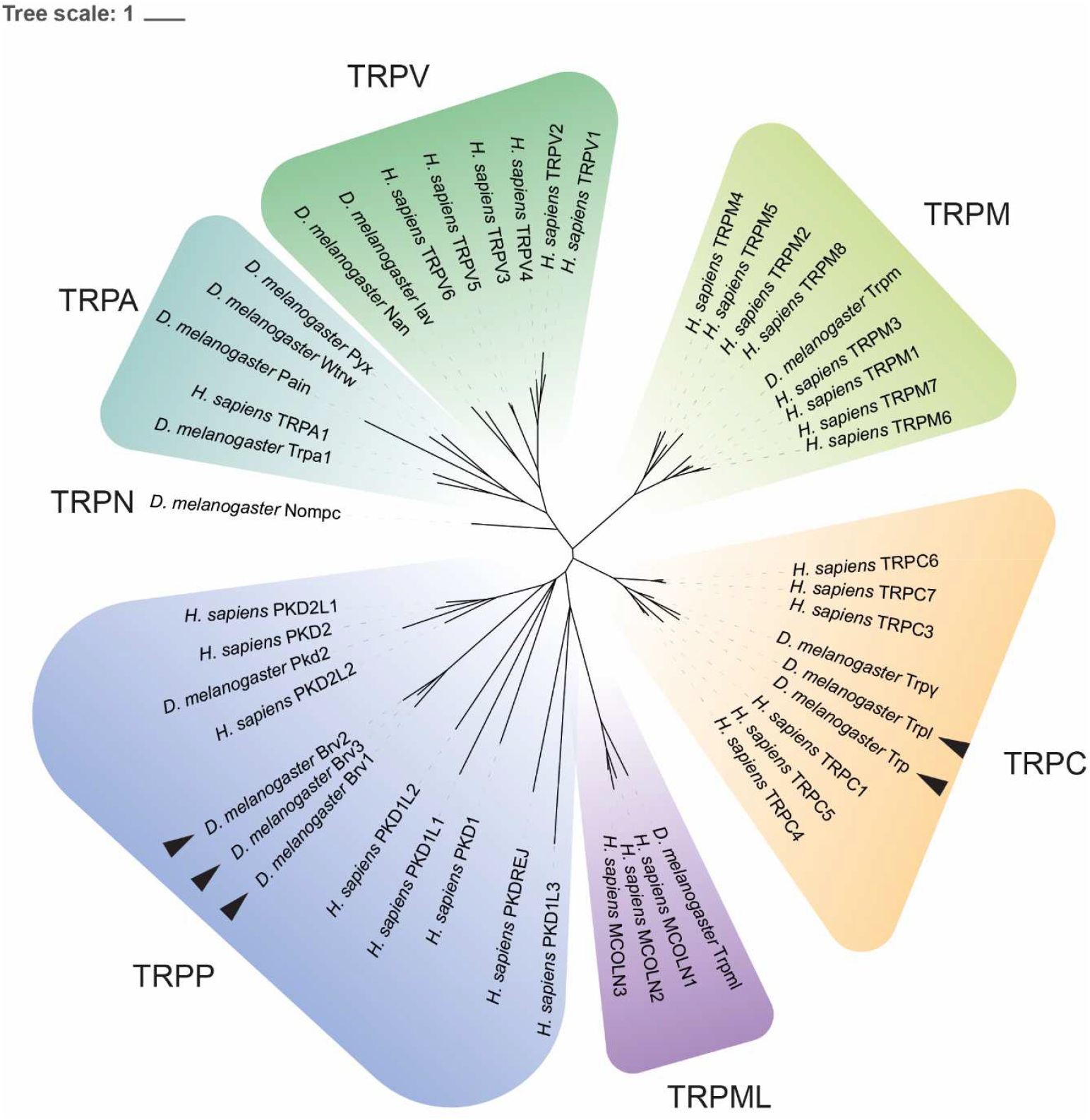
Phylogenetic tree reconstruction of *Drosophila* and human TRP channels highlighting the different classes within this protein family. Black arrows mark TRP channels analysed in the present study. For bootstrap values, please refer to Supplementary Figure S1.

Cold temperatures are perceived through distinct sensory structures, and through different mechanisms in *Drosophila* larvae and adults. In larvae, chordotonal organs (COs), the terminal organ (TO) and Dorsal Organ Cool Cells (DOCCs) represent the main structures mediating cold sensing, with both TRP channels and Ionotropic receptors (IRs) playing a role. Mutations affecting *transient receptor potential* (*trp*) and *transient receptor potential-like* (*trpl*) loci lead to impaired avoidance of temperatures as low as 10°C, suggesting that these TRP channels, which are expressed in the TO, control larval cool avoidance [32]. In adult flies, Trp and Trpl are relevant to phototransduction in the eyes through the phospholipase C encoded by *no receptor potential A* (*norpA*) [32], with no reported roles in thermosensation. In line with this, inhibiting the activity of *GH86-Gal4* expressing neurons (TO) in larvae results in impaired cool avoidance but does not affect the response to warm temperatures, suggesting that warm and cool avoidance depend on distinct neuronal circuits [32]. By contrast, impaired firing of *GH86*-expressing neurons in adult flies has been shown to exclusively impact warm avoidance [33], possibly suggesting neuronal network remodelling during development. Similarly, mutations affecting *inactive* (*iav*), a TRP channel specifically expressed in COs, prevent larvae from choosing their preferred temperature (17.5°C) over slightly cooler ones (14-16°C) [34]. More recent studies have implicated also ionotropic receptors Ir21a, Ir25a and Ir93a - which are located in the DOCCs - in larval cool avoidance, as their respective null mutants, when placed in the middle of a thermal gradient ranging from 13.5°C to 21.5°C, show impaired ability to avoid colder temperatures [35,36]. On the other hand, different TRP channels expressed in larval cold-sensing neurons, such as Pkd2, NompC and Trpm, seem to mediate *Drosophila* responses to noxious cold (≤10°C) [37].

In adult flies the main thermosensory structures are located in the antennae. In 2011, Gallio et al. [15] identified three novel TRP channels, called Brivido (Brv) 1, 2, and 3, expressed in the arista and sacculus. *brv* knockouts (*brv1* and *2*) and knockdown (*brv3*) display significant defects in cool avoidance [15], at temperatures as low as 11°C, thus within the range of those triggering dormancy. Another study suggests that the perception of cool innocuous temperatures (16°C) – but also warm ones (31°C) – is mediated by Ir21a, Ir25a and Ir93a ionotropic receptors located in Aristal Cold/Cooling Cells, and does not require Brv TRP channels [38]. While all the above evidence suggests that cold-sensing relies on different molecular machinery in larvae and adult flies, Turner et al. have recently reported that chordotonal neurons and *brv1* mediate cold nociceptive sensitization (5-15 °C) following UV-induced tissue damage in larvae [39], thus pointing to the existence of cold-sensing mechanisms common to different developmental stages. Overall, the majority of the literature on both mammalian and *Drosophila* temperature perception suggests that different molecular mechanisms are responsible for the perception of cool, cold and noxious cold temperatures.

Similarly to mutations affecting temperature perception, also impaired brain neurotransmitter and intracellular pathways have been shown to affect temperature preference in flies. Specifically, mutants of *adenylyl cyclase* (*rutabaga, rut*^*1*^) and *cAMP phosphodiesterase* (*dunce, dnc*^*1*^) exhibit impaired avoidance of cold and warm temperature, respectively, suggesting a direct relationship between cAMP levels (particularly in the mushroom bodies) and preferred temperature [33]. Moreover, flies carrying mutations in genes encoding enzymes/receptors implicated in dopamine and histamine synthesis/signalling show an overall preference for colder and warmer temperatures, respectively [40,41].

Most cold sensing studies in both mammals and insects have investigated relatively simple behavioural responses such as thermotactic behaviour/cool avoidance, with less attention being devoted to more complex processes such as growth and reproduction. Similarly, and despite the large body of literature on the molecular and neuronal mechanisms underlying cold-sensing and temperature preference, only few studies have focussed on their contribution in orchestrating growth/developmental adaptive processes (see [42–44]), such as overwintering strategies triggered by a decrease in temperature (i.e. dormancy).

Here we use ovarian development at low temperature as a case study, utilising *Drosophila* genetics to investigate the contribution of cold-sensing TRP channels in the induction of reproductive dormancy, a primarily cold-induced arrest/slowing of gonads maturation [45–48]. More specifically, we present functional phenotypic evidence of the involvement of the cold-sensing TRP channels Trp, Trpl and Brv1 in orchestrating gonadal slowing/arrest at low temperature in *Drosophila*. Further, we provide evidence of the relevance of temperature preference for tissue/organ growth under mildly stressful conditions like those triggering dormancy.

## 2. Results

### 2.1 Drosophila mutants for the cold-sensing TRP channel brivido 1 exhibit enhanced ovarian development at low temperatures

We tested whether impaired cold-sensing signalling in adult flies is sufficient to antagonize ovarian dormancy. To this aim, mutants for the *brv1* gene, which mediates perception and avoidance of a range of temperatures including those triggering dormancy (11-14°C) [45,48], were assayed. Using an established experimental paradigm [43,45,49], both hypomorphic (NP4486 called *brv1*^*hyp*^) and loss-of-function (*brv1*^*L563 > STOP*^ called *brv1*^*-/-*^) *brv1* mutant females exposed at 12°C and light-dark (LD) cycle 8:16 were found to exhibit significantly enhanced ovarian development compared to their controls (p < 0.0001; Figure 2A). The ovaries of *brv1*^*-/-*^ females appeared highly vitellogenic, including stage 14 oocytes (the last stage that is then laid), as well as exhibiting a decrease in ovarioles number (Figure 2D, E) compared to control flies (Figure 2F).

**Figure 2.**
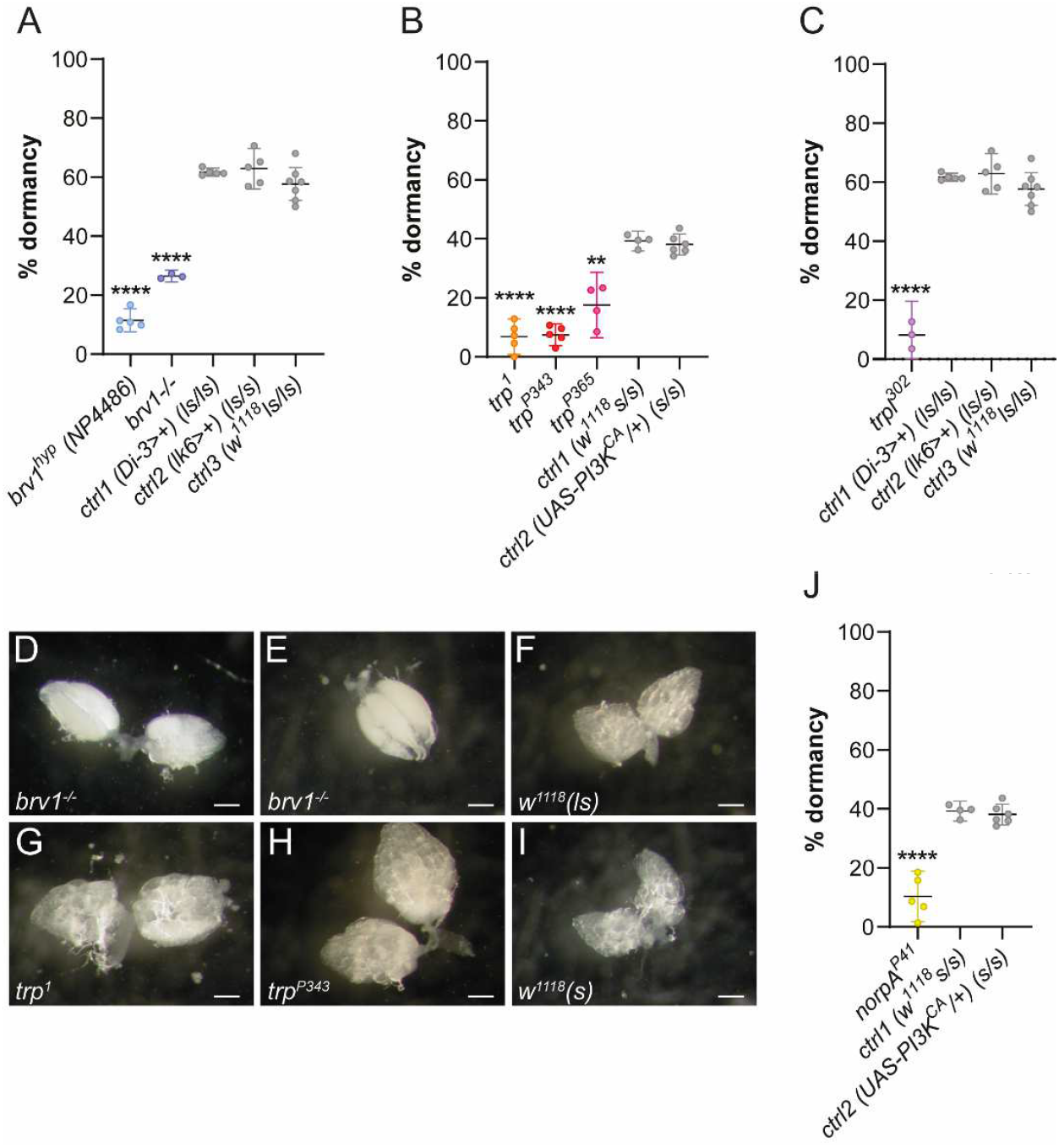
Percentage ovarian dormancy (mean ± 95% confidence interval) in *brv1* mutants (A) (*brv*^*hyp*^ and *brv1*^*-/-*^) and respective background controls; significance (**** p < 0.0001) refers to post hoc ANOVA comparisons of arcsine square-rooted data. Percentage ovarian dormancy (mean ± 95% confidence interval) in females of (B) *trp* (*trp*^*1*^, *trp*^*P343*^, *trp*^*P365*^) and (C) *trpl* mutants, and pertinent background controls. Controls used in (A) and (C) are the same, as experiments were run in parallel; significance (** p < 0.01, **** p < 0.0001) refers to post hoc ANOVA comparisons of arcsine square-rooted data. Representative pictures of ovarian development after 11 days at 12°C in *brv1* (D-E) and *trp* (G-H) mutants, with pertinent controls, i.e. *w*^*1118*^ (*ls-tim/ls-tim*) (F) and *w*^*1118*^ (*s-tim/s-tim*) (I). White bars = 0.2 mm. Percentage (mean ± 95% confidence interval) of dormant females in *norpA*^*P41*^ mutants (J) and pertinent background controls. Controls in (J) are the same as in (B), as experiments were run in parallel; significance (**** p < 0.0001) refers to post hoc ANOVA comparisons of arcsine square-rooted data.

### 2.2 Loss-of-function mutations at the trp and trpl genes promote ovarian development at low temperatures

Additional, loss-of-function mutants for the genes *trp* and *trpl* encoding TRP channels were tested. These mutants have already been shown to modulate perception and avoidance of temperatures as low as 10°C in behavioural assays [32]. Three independently generated *trp* mutants (*trp*^*1*^, *trp*^*P343*^, *trp*^*P365*^) showed a significant increase in ovarian growth compared to pertinent controls (** p < 0.01 and **** p < 0.0001, respectively; Figure 2B). In these mutants, ovaries were larger and more developed (Figure 2G, H) compared to those of controls (Figure 2I), with advanced vitellogenesis, similarly to *brv1*^*-/-*^ mutants (Figure 2D-E). Similarly, loss of function mutants for a third TRP channel, *trpl* (*trpl*^*302*^) also exhibited significantly decreased levels of ovarian dormancy (p < 0.0001; Figure 2C). Finally, at low temperatures, ovarian development was significantly enhanced in *norpA*^*P41*^ null allele mutants, similarly to what observed for *trp* and *trpl* mutants (p < 0.0001; Figure 2J).

### 2.3 Mutations modulating temperature preference affect ovarian development at low temperatures

To test the hypothesis that preference for lower or higher temperatures than those preferred by wild-type flies (∼24°C; [33,40,50]) could affect growth/developmental/reproductive trajectories, the incidence of ovarian dormancy was measured in mutants characterized by shifts in temperature preference [i.e. for the genes *rutabaga* (*rut*), *dunce* (*dnc*), and *ora transientless* (*ort*)]. More specifically, we predicted that mutants with impaired cold avoidance (*rut*^*1*^) would show more pronounced ovarian growth in the cold, while mutants with impaired warm avoidance (*dnc*^*1*^) would show dampened ovarian development in the cold. Unlike *rut*^*1*^ and *dnc*^*1*^ mutants, *ort*^*1*^ mutants [with *ort* encoding an ionotropic histamine-gated chloride channel required for photoreceptor signalling and temperature preference settings in the central brain [41]], show a more complex temperature preference, with reduced avoidance of both colder and warmer temperature compared to *w*^*1118*^ controls, with a shift towards slightly warmer temperatures [41]. Therefore, we opted to test *ort*^*1*^ under both short and long photoperiods (please also refer to Materials and methods). We found *rut*^*1*^ and *ort*^*1*^ loss-of-function mutations to be associated with a significant decrease in ovarian dormancy, even under short photoperiods (LD 8:16) (Figure 3A). By contrast, *dnc*^*1*^ mutants showed an increase in ovarian dormancy under long photoperiods (LD 16:8) (Figure 3B). Conversely, *ort*^*1*^ females maintained low dormancy levels also under long photoperiods (LD 16:8) (p < 0.0001; Figure 3B).

**Figure 3.**
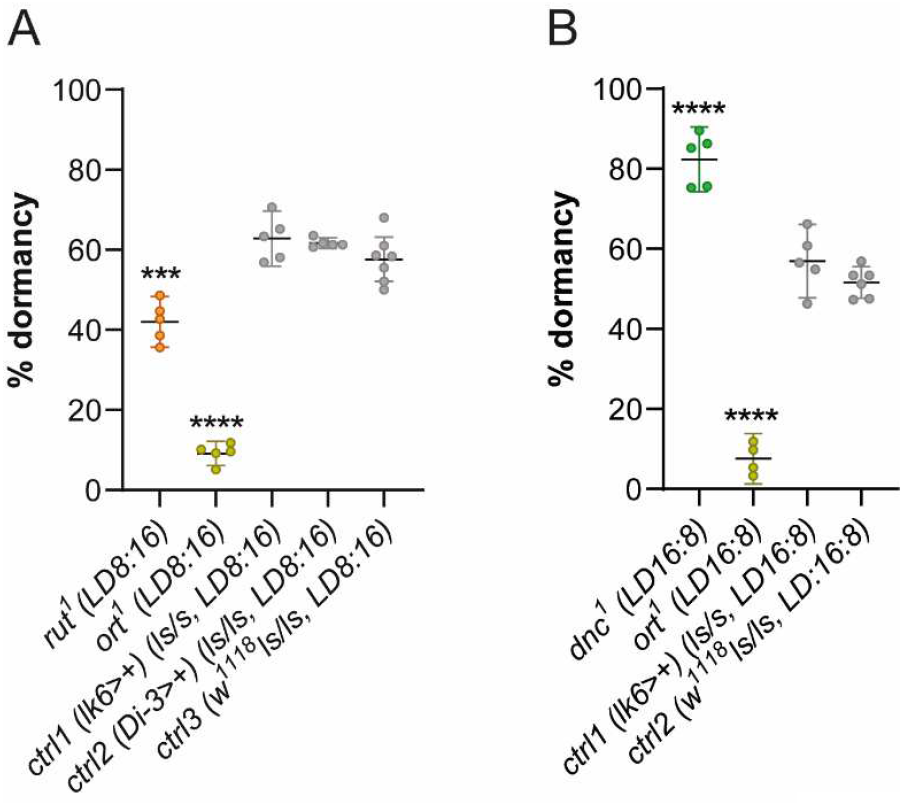
Percentage ovarian dormancy (mean ± 95% confidence interval) in *rut*^*1*^ and *ort*^1^ mutants (A) and pertinent background controls. Controls used in (A) are the same as in Figure 1A and C, as experiments were run in parallel; significance (*** p < 0.001) refers to post hoc ANOVA comparisons of arcsine square-rooted data. Percentage ovarian dormancy (mean ± 95% confidence interval) in *dnc*^*1*^ and *ort*^*1*^ mutants (B) and pertinent background controls. Ctrl2 are the same as in [43], as experiments were run in parallel; significance (**** p < 0.0001) refers to post hoc ANOVA comparisons of arcsine square-rooted data.

## 3. Discussion

The role of temperature in modulating ovarian development in *D. melanogaster* is well established [43,45,49,51,52] but the exact mechanisms through which it triggers ovarian dormancy remain unclear. One possibility is that dormancy might be triggered by brain thermosensory cascades modulating neuroendocrine pathways for growth, development and reproduction [42,43], as already known for behavioural responses [15,32,33,40,41]. Alternatively, the phenomenon might occur as a consequence of cold-induced systemic physiological changes, such as a drop in enzymatic activity and a slowing in metabolic rates, as part of a stress response [46]. This would, in turn, affect gonadal maturation/dormancy directly, or as a consequence of feedback from the periphery to the brain [53]. This study provides the first functional evidence of the involvement of cold-sensing TRP channels in modulating ovarian development. More specifically, we show that loss-of-function mutants for TRP channels involved in the perception of non-noxious cold reduce dormancy levels.

Surprisingly, our data show that the *brv1*^*hyp*^ hypomorphic allele is more effective in enhancing ovarian growth at low temperature compared to the *brv1*^*-/-*^ loss of function. The transposable element inserted in the hypomorphic strain acts as a Gal4 enhancer trap (*NP4486-Gal4* or *brv1-Gal4*, see [15,35,54]), and it is located ∼2kb downstream of the *brv1* gene and 2.5kb upstream of a non-coding RNA gene (CR32207, [35,54]), which has been recently linked to some features of oogenesis [55]. Different lines of evidence clearly suggest that both *brv1* and *CR32207* are expressed in the adult antennae [15,35,56,57], in accordance with the expression of the *NP4486-Gal4* construct, which is detected in the entire structure. However, Klein et al. identified *brv1-Gal4* expression in both cool-sensing neurons as well as other sensory neurons in larvae [54]. It is therefore possible that the stronger effects exerted by *brv1*^*hyp*^ on ovarian dormancy may be due to the involvement of *CR32207*, and consequent impairment in specific classes of sensory neurons. On the other hand, *brv1*^*-/-*^ females exhibited higher vitellogenic levels in eggs, despite the clearly reduced number of ovarioles, suggesting a more robust effect of this loss-of-function mutation on ovarian development at 12°C compared to *brv1*^*hyp*^ flies.

We also provide evidence for the involvement of Trp and Trpl in dormancy induction/maintenance, with mutants for these genes exhibiting highly vitellogenic well-developed ovaries at low temperatures. Interestingly, mutations in these genes have also been associated with significant deficits in larval cool avoidance [32]. In larvae, the expression of *trp* and *trpl* has been reported in the TO, one of the main cold-sensing structures at this developmental stage. However, in adult flies these TRP channels are mostly known for their role in phototransduction [58,59]. To date, with *trp* and *trpl* being primarily expressed in the visual system [58,59], no studies have pointed to a potential dual photo-/thermo-sensory role for these genes. Interestingly, Shen et al. documented a thermosensory role for rhodopsin, a rhabdomeric photoreceptor in *Drosophila* eyes and homologous to human OPSD/Rhodopsin [60]. In its temperature-sensing role, rhodopsin is coupled with a downstream TRP channel, dTRPA1 [60]. Further, various forms of metabolic stress (i.e. anoxia, mitochondrial uncouplers, and ATP depletion) have been shown to activate Trp and Trpl *in vivo* [61], supporting the hypothesis that cold-induced metabolic slowing - as opposed to cold perception *per se* - may trigger a similar mechanism. Of note, one of the *trp* mutant strains used in this study, *trp*^*1*^, has been reported to carry a temperature-sensitive allele inducing altered cold avoidance when flies are reared at moderately warm temperatures (25-27°C) [32]. Taken together, our data on *trp* and *trpl* knockouts point to a role of these TRP channels in adult thermosensation for moderate cold, similarly to what has previously been shown in larvae [32]. Considering the well-known role of Trp and Trpl in phototransduction, a light-dependent effect on dormancy propensity mediated by these TRP channels could also contribute to our findings. Brv1, Trp and Trpl mammalian homologs (TRPP and TRPC family members, respectively, see Figure 1; Supplementary Figure S1) have also been shown to play important roles in regulating developmental features and reproduction [62–69], temperature sensing and environmental sensing at large [70–72]. Concerning the downstream signalling pathways, the involvement of NorpA (ortholog of human Phospholipases C β4, see Supplementary Figure S2) in the thermosensory cascade seems negligible, at least in larvae [32]. This is in contrast with our observations in adults, where this phospholipase mediates phototransduction [73].

We also provide evidence of the influence of pathways controlling temperature preference on complex developmental trajectories in *Drosophila*. Altered ovarian development was observed at low temperature in *rut, dnc* and *ort* mutants, with the direction of the effects being in line with our predictions (based on the preference shift towards cooler and warmer temperatures in *rut*^*1*^ and *dnc*^*1*^ mutants, respectively). By contrast and despite their impaired warm avoidance, *ort*^*1*^ mutants exhibited reduced ovarian dormancy irrespective of photoperiod, potentially implicating histamine signalling [as well as dopamine, serotonin and octopamine signalling [43]] in dormancy-induction. Indeed, we have previously shown [43] reduced dormancy levels in mutants of genes encoding important components of dopamine synthesis/signalling (i.e. *ple*^*4*^, *ddc*^*DE1*^, *dop1r1*^*hyp*^), which are characterized by impaired cold avoidance [40].

Rut, Ort and Dnc are the fly orthologs of human ADCY1, GLRA1-3 and Phosphodiesterases PDE4A-D, respectively (Supplementary Figures S3, S4, S5). As Rut and Dnc enzymes control cAMP signalling, which is involved in a variety of physiological, endocrine and metabolic functions, it is not surprising to find their mammalian orthologs associated with the development of the female reproductive system and fertility [74–77]. Further, inhibitors of PDE4 have been shown to induce hypothermia in mice [78], and to abolish stress response - again, triggered by hypothermia - mediated by Cold-Inducible RNA-Binding Protein [79].

Responses to variations in environmental temperature are crucial for survival. This is why animals exhibit a temperature preference, which is particularly robust in poikilotherms. Thus, coherent changes in dormancy propensity in mutant strains characterized by impaired thermal preference are not unexpected, as such impairment impinges on physiology, metabolism, energy allocation, and ultimately developmental/reproductive trajectories, depending on the temperature animals are exposed to. The manipulation of such reproductive trajectories may also be relevant to the development of novel strategies for pest insect species proliferation control [80]. Further, genes like *dnc, rut, ort* are not exclusively involved in setting temperature preference but have pleiotropic functions, that might directly or indirectly affect dormancy propensity. For instance, *dnc* is required for *Drosophila* oogenesis [81], and *ort* has been found to be expressed in testis [82], but not in the eggs [83].

As a whole, our phenotypic analysis provides functional support for a role of cold-sensing pathways and temperature preference in modulating developmental and reproductive processes.

The hypothesis that temperature preference might directly or indirectly impinge on species-specific developmental and reproductive features is supported by several lines of evidence. For instance, in the Nile tilapia and the African catfish, sex determination is influenced by the temperature experienced early in development [84,85]. Similarly, migratory locusts have been shown to prefer temperatures where growth rate increases despite a less efficient nutrient utilization [86]. Ultimately, temperature preference has an adaptive value, as nicely demonstrated by larvae/flies from high and low latitudes preferring cooler and warmer temperatures, respectively [87].

In humans, loss- or gain-of function mutations of TRPs cause a number of inherited diseases called channelopathies [19]. For example, TRPV4 channelopathy is linked to at least nine different diseases, ranging from autosomal dominant brachyolmia type 3 to Charcot-Marie-Tooth neuropathy type 2 [88]. Further, polymorphisms in TRP genes may regulate disease risk, for example with carriers of rs10166942, which correlates to reduced TRPM8 gene expression, having a lower incidence of migraine [89]. Similarly, a single nucleotide polymorphism in the TRPC6 gene has been identified in patients with the autoimmune disease lupus erythematosus [90]. Up- or down-regulation of TRPs has been associated with a large and diverse set of human disorders, which is not surprising given that TRPs are multifunctional signalling molecules with multiple roles in sensory perception and cellular physiology [91]. Accordingly, they have also been proposed as therapeutic targets for diseases as diverse as metabolic, cardiovascular and neurological, and certain steps of cancer progression [19,91]. Drug discovery efforts to target TRP channels originally focused on pain. More recent applications in mammalian models include, for example, Parkinson’s disease. Here, an antibody against the heat-sensitive TRPV1 - within a photothermal wireless deep brain stimulation nanosystem - has been shown to counteract dopamine neurons degeneration [92]. Nano-delivery systems encompassing metal nanostructures to generate heat and thus modulate thermally activated TRP channels are of more general interest, as they can improve targeting of existing drugs [20]. A recent human single-arm, open label multicentre study documented that an inhibitor of TRPV2 expression was effective in preventing cardiac function worsening in muscular dystrophy patients [93]. Finally, ongoing human trials include the assessment of TRP-targeting compounds for chemotherapy-induced peripheral neuropathy, dysphagia and aging (https://clinicaltrials.gov/).

The most obvious mammalian correlates of our findings may relate to metabolism [20] and development. Of interest and as an example, TRPC5 has been implicated in the central regulation of energy and glucose homeostasis [94]. More generally, TRPs mediate the recruitment of brown adipose tissue both via cold exposure and as chemesthetic receptors for certain food ingredients, thus impinging on energy dissipation and weight [95]. Finally, while metabolism and metabolic rates have obvious effects on development, it is also of interest that TRPs, including those of the classes we studied here, are expressed in both the ovary and testis [20]. Therefore, the hypothesis that TRPs may modulate gonadal development and development at large in mammals appears plausible.

## 4. Materials and methods

### 4.1 Fly stocks and maintenance

Fly stocks were maintained at 23 °C in a 12:12 hour light/dark (LD) cycle prior to the experiments. For purposes of both stock maintenance and the experiments described here below, a standard yeast-sucrose-cornmeal diet was used [43]. The following fly strains were used: *NP4486* (called *brv1*^*hyp*^) and *brv1*^*-/-*^ (provided by Marco Gallio, Northwestern University, and Charles S. Zucker, Columbia University), *norpA*^*P41*^ (provided by Charalambos Kyriacou, University of Leicester), *hmgcr*^*Di-3*^*-Gal4* (provided by Jean-René Martin, University of Paris-Saclay), *w*^*1118*^*(s-tim)* (provided by Charlotte Helfrich-Foster, University of Würzburg), *w*^*1118*^*(ls-tim). trp*^*1*^ (5692), *trp*^*P343*^ (9046), *trp*^*P365*^(9044), *trpl*^*302*^(31433), *rut*^*1*^ (9404), *dnc*^*1*^ (6020), *ort*^*1*^ (1133), *UAS-PI3K*^*CAAX*^ (called *UAS-PI3K*^*CA*^, 8294), and *Lk6*^*DJ634*^*-Gal4* (8614) were obtained via the Bloomington Drosophila Stock Center. The *brv1*^*hyp*^ hypomorphic allele was generated with the insertion of a transposable element (*P{GawB}*) 2249 bp downstream of the *brv1* stop codon [15]. The *brv1*^*-/-*^ (*brv1L563STOP*) is an amorphic allele bearing a nucleotide substitution (T1683A), which truncates the native protein within the highly conserved ion transporter domain [15]. *trp*^*1*^, *trp*^*P343*^, *trp*^*P365*^, and *trpl*^*302*^ are loss of function alleles of *trp* and *trpl* genes (see [58,59,96–98]). *trp*^*1*^ is a temperature-sensitive allele which shows altered cold avoidance when reared at 25-27°C but not at 18°C [32]. The *norpA*^*P41*^ allele bears a 351 bp deletion which causes a frame-shift and results in the substitution of 120 amino acids and a premature stop codon within the catalytic domain. Further, the resulting protein lacks the C-terminal required for Gαq interaction [99].

### 4.2 Genetic controls and genetic background

Polymorphisms at specific loci (i.e. *timeless* [*tim*] and *couch potato* [*cpo*]) have been shown to affect the propensity of *Drosophila* females to enter/maintain reproductive dormancy [100–102]. Further, we have previously demonstrated, in experimental conditions similar to those of the present study [43,45], that the effects of the polymorphism at the *tim* locus (with the two variants named *s-tim* and *ls-tim*, [100,103]) are stronger than those associated with *cpo* gene polymorphisms [101,104]. Therefore, here we focussed our genotyping strategy exclusively on the *s-tim* - *ls-tim* polymorphism. Genotyping of the *tim* locus was performed as described in [43,100]. Briefly, genomic DNA was extracted from 5-10 females from each strain, and amplified using Amplification Refractory Mutation System (ARMS) PCR [100], with the following primers: *ls-tim* forward: 5’-TGGAATAATCAGAACTTTGA-3’; *s-tim* forward: 5’-TGGAATAATCAGAACTTTAT-3’; *s-tim* reverse: 5’-AGATTCCACAAGATCGTGTT-3’ (common).

As described in [43,45,49], for the experiments involving mutant strains, we used *w*^*1118*^ (*s-tim* or *ls-tim* based on the mutant background), and GAL4 or UAS lines (of appropriate *tim* background) crossed to *w*^1118^ as controls. The controls’ background at the *tim* locus is specified in each relevant figure.

### 4.3 Reproductive dormancy assays

Levels of reproductive dormancy were assessed using a previously published protocol [43,45,49]. Briefly, larvae were reared under standard conditions at 23 °C and LD 12:12 until eclosion. Newly eclosed virgin flies were collected (∼60 females and 60 males *per* replicate, unless otherwise specified) within 5 hours of eclosion, and rapidly exposed to low temperatures (12 °C) and short (LD 8:16) or long (LD 16:8) photoperiods for 11 days. Similarly to previous study of ours [43,45], with mutations expected to increase ovarian development at 12°C, we assessed dormancy under LD 8:16 to further strengthen dormancy-inducing conditions. Conversely and based on the same principle, when mutations were expected to reduce ovarian development flies were tested under LD 16:8. Unless otherwise specified, all strains were tested under LD 8:16. Reproductive dormancy was defined as the complete absence of vitellogenesis (i.e. all oocytes at stages ≤ 7), examining all ovarioles in both ovaries of each specimen [45,48,100] on day 11. Three-seven biological replicates (n ≥ 46 females each, ∼180-420 flies in total) were analysed for each genotype, with the exception of *brv1*^*-/-*^ for which a total of ∼120 females were available (3 biological replicates of n ≥ 35 females each). Data on *w*^*1118*^*(s-tim)* and *w*^*1118*^*(ls-tim)* are the same used in [43], as the experiments described in [38] and those described here were performed at the same time. Dormancy levels are presented as percentage of dormant females. Percentage data were arcsine square-root transformed and analysed by one-way ANOVA (*post hoc* Tukey test) [43,45,100] using GraphPad Prism 9.0.0.

### 4.4 Phylogenetic study and sequence analysis

To reconstruct phylogenetic relationships for the proteins encoded by the genes of interest (*brv1, trp, trpl, norpA, rut, dnc, ort*), their aminoacidic sequence predicted by Himmel and Cox [26] were utilized (FlyBase or NCBI databases). Protein sequences from species other than *D. melanogaster* were retrieved by Basic Local Alignment Search Tool (BLAST) from NCBI databases. Hit sequences were aligned using MUSCLE [105]. Maximum likelihood phylogenies were generated using the IQ-TREE [106] web server, with default settings. Consensus trees generated were visualized using the Interactive Tree of Life tool [107] and rooted, where possible, using a protein outgroup identified by BLAST searches [108]. Protein sequences used for phylogenetic reconstructions and their respective accession numbers are presented in Supplementary Table S1.

## Data availability statement

The raw data supporting the conclusions of this article will be made available by the corresponding authors upon reasonable request.

## Author contributions

Concept and design, GA and RC; experiments, GA; data analysis and interpretation, GA, SM and RC; writing of the manuscript, GA; review of the article for important intellectual content, SM and RC; funding acquisition, RC and SM.

