## Supplementary figures and images for "Cold-sensing TRP channels and temperature preference modulate ovarian development in the model organism *Drosophila melanogaster*"

### Supplementary Figure 1

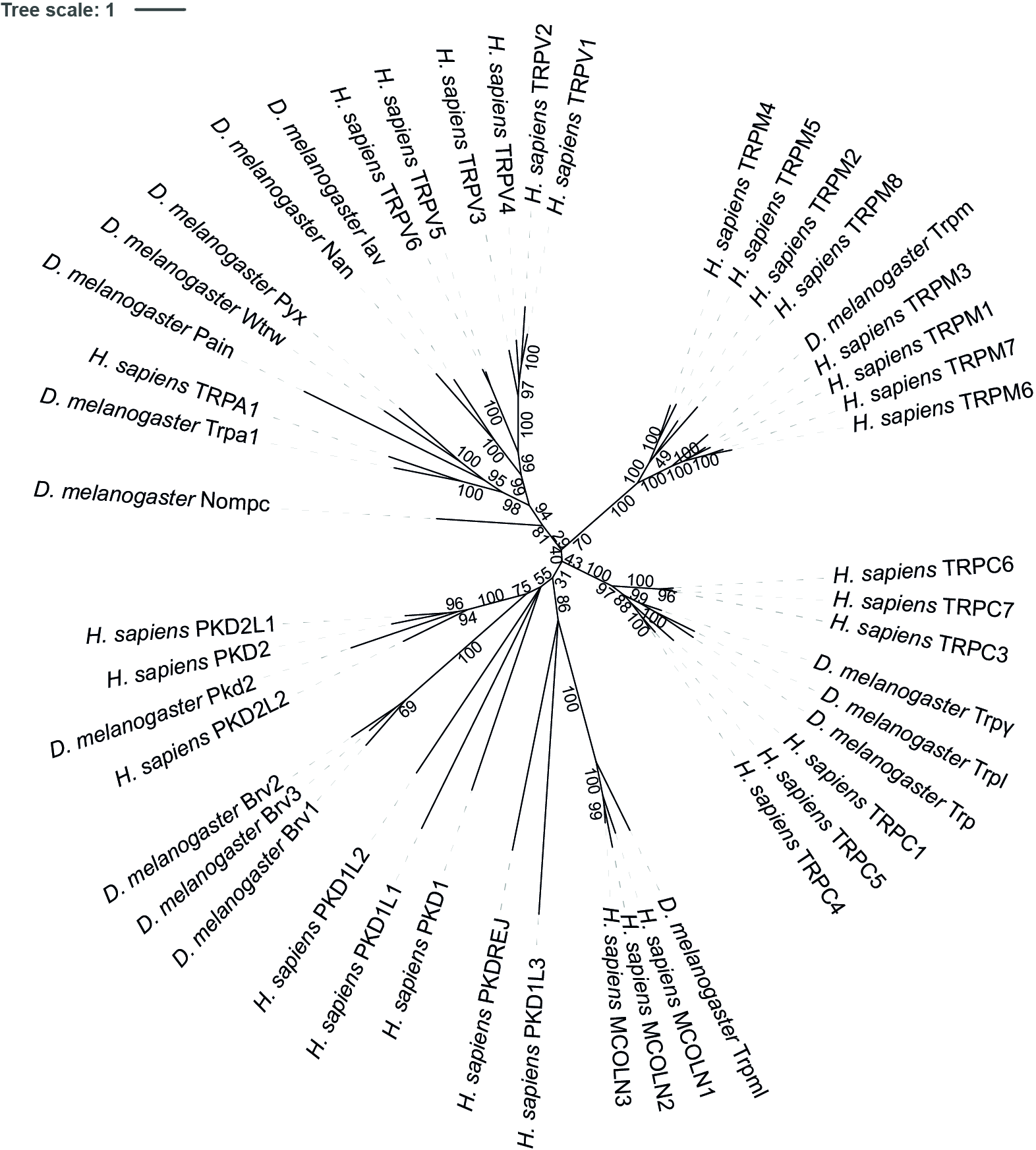

### Supplementary Figure 2

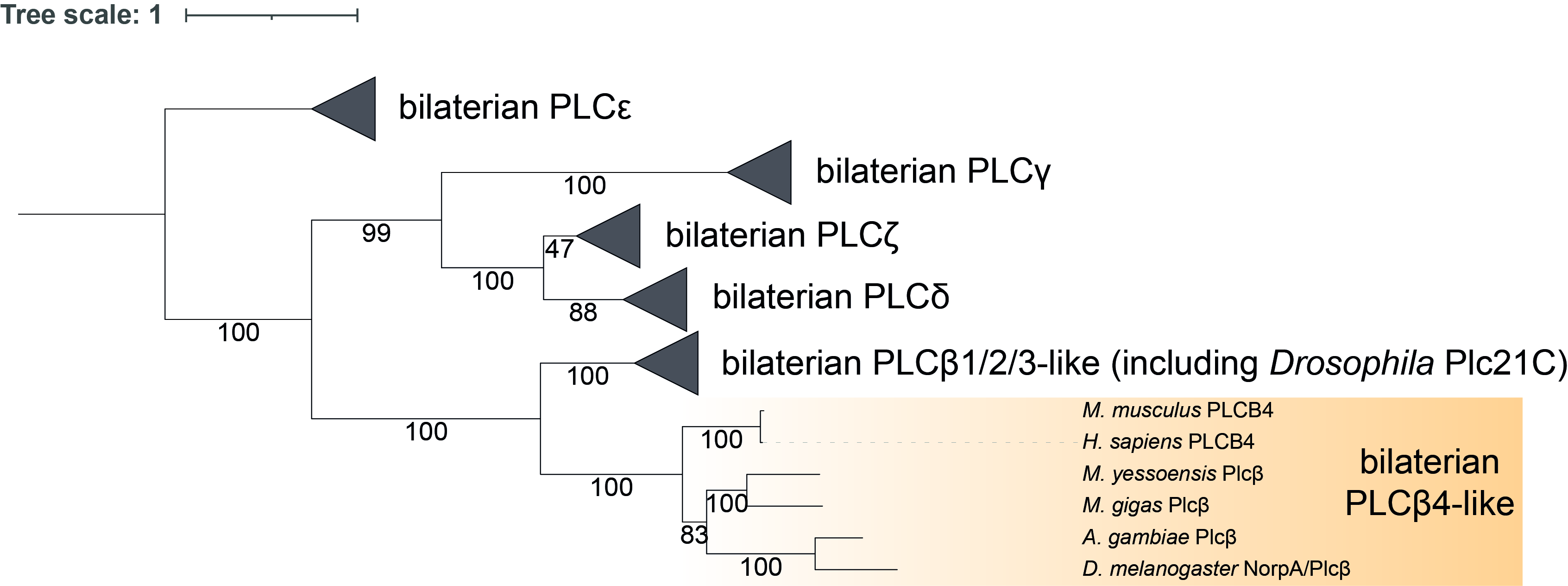

### Supplementary Figure 3

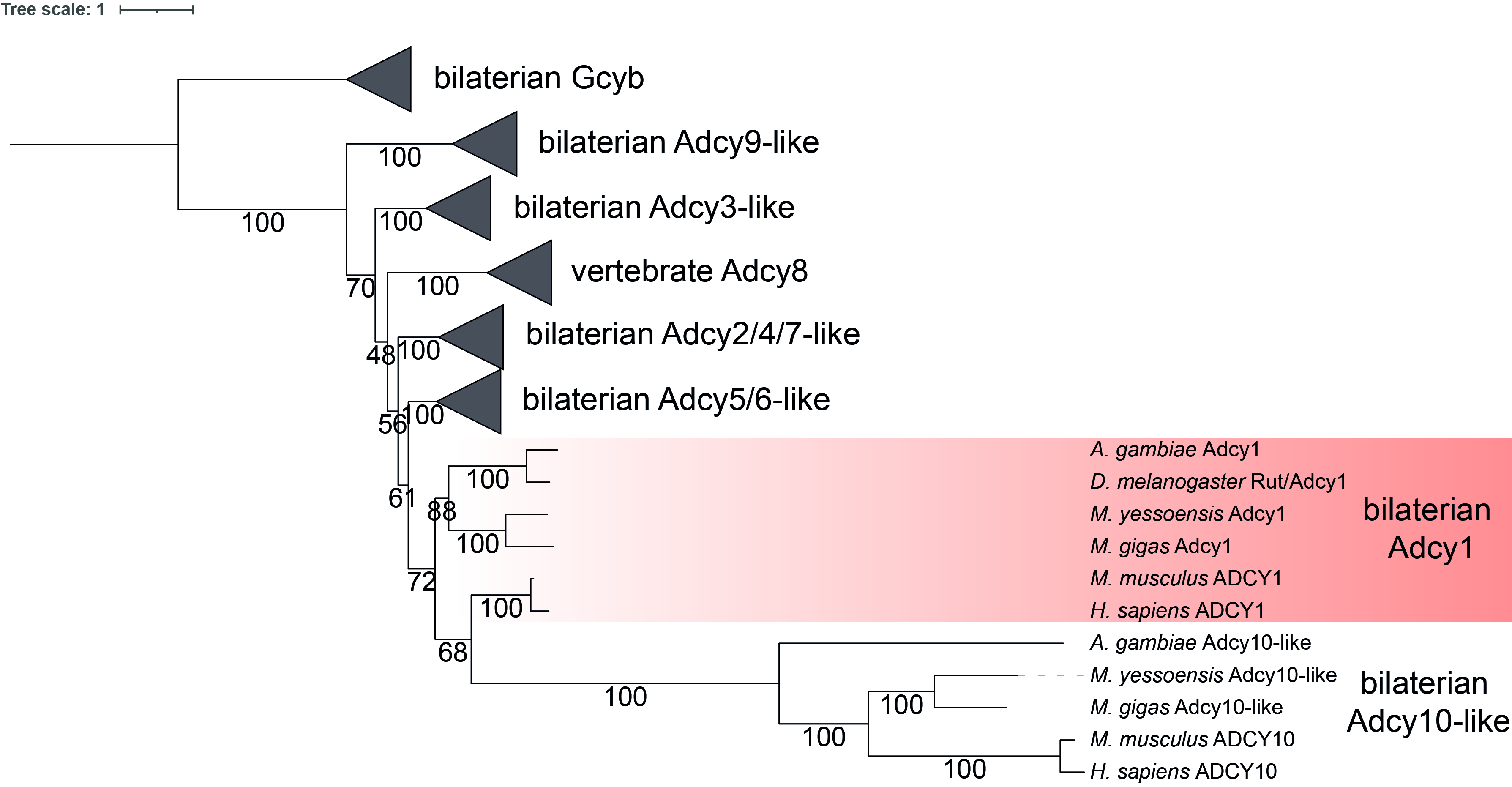

### Supplementary Figure 4

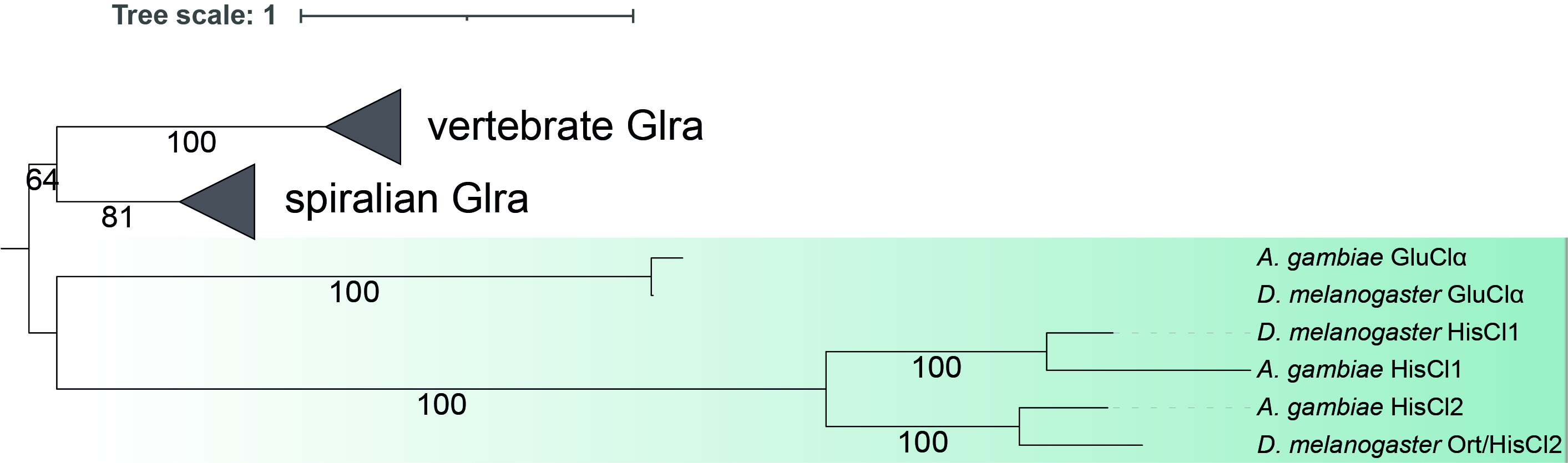

### Supplementary Figure 5

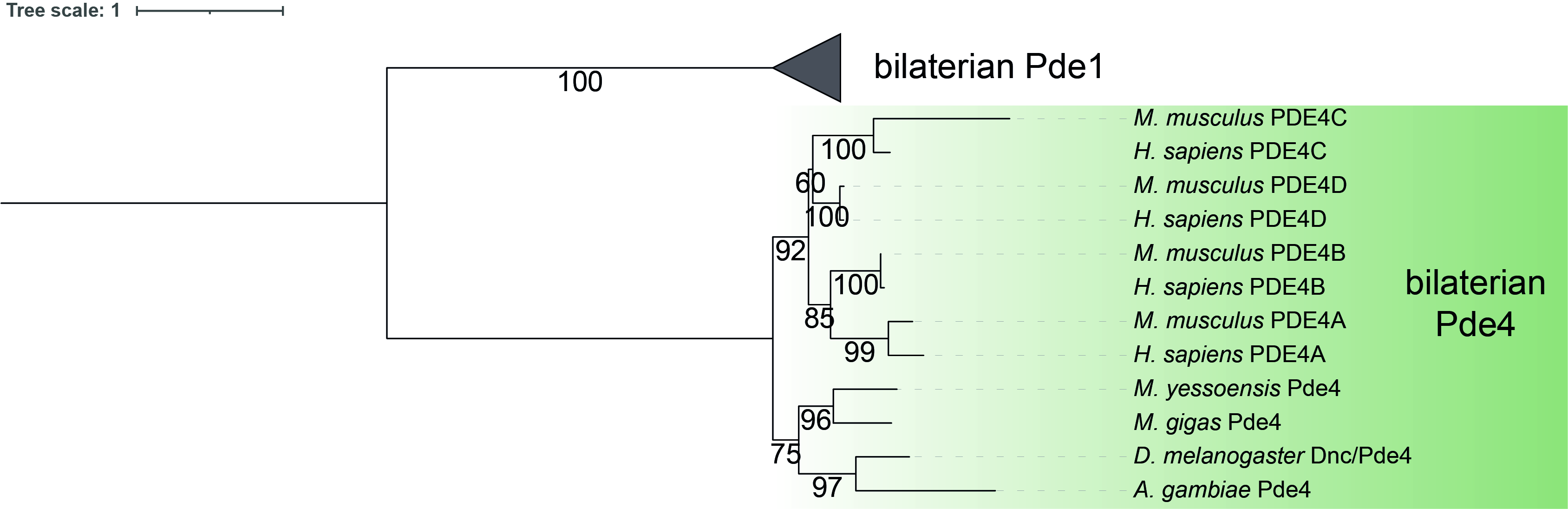
